# Cold acclimation of *Trogoderma granarium* Everts is tightly linked to regulation of enzyme activity, energy content and ion concentration

**DOI:** 10.1101/296467

**Authors:** Mozhgan Mohammadzadeh, Hamzeh Izadi

**Affiliations:** Department of Plant Protection, Faculty of Agriculture, Vali-e-Asr University of Rafsanjan, Rafsanjan, Iran

**Keywords:** Cold acclimation, Fluctuated thermal treatment, Rapid cold hardiness, The Khapra beetle

## Abstract

In this study, cold hardiness and some physiological characteristics of *T. granarium* larvae were investigated under different thermal regimes, i.e. warm-acclimated (WA), cold-acclimated (CA), fluctuating-acclimated (FA) and rapid cold-hardened (RCH). In all regimes, the survival rate of the larvae decreased with a decrease in temperature and raise in exposure time. Cold acclimated larvae showed the highest cold hardiness in -15 and -20 ºC. Control larvae had the highest glycogen content (34.4 ± 2.3 µg/gdw). In contrast, cold acclimation larvae had the lowest glycogen content (23.0 ± 1.6 µg/gdw). Change in trehalose content was reversely proportional to change in glycogen content. The greatest myo-inositol and glucose contents were detected in larvae cold acclimation treatment (10.7 ± 0.4 µg/gdw) and control (0.49 ± 0.03 µg/gdw), respectively. In control and treated larvae, the concentration of Na^+^ decreased, though the concentration of K^+^ rose, with rising the exposure time. The shape of the thermal reaction of AMP-depended protein kinase and protein phosphatase IIC followed the same norm, which is different from protein phosphatase I and protein phosphatase IIA. Protein phosphatase IIA and IIC showed a complete difference in thermal reaction norms. In did, thermal fluctuation caused the highest changes in the activity of the enzymes, whereas the RCH showed the lowest changes in the activity of the enzyme. Our results showed a significant enhancement of larval cold tolerance under CA regime that is related to the level of low molecular weight carbohydrates, protein kinase, and phosphatases activity, and hemolymph ions concentration.

**SUMMARY STATEMENT:** In *Trogoderma granarium*, cold acclimation enhances the larval cold tolerance that is related to change in the level of low molecular weight carbohydrates, protein kinase, and phosphatases activity, and hemolymph ions concentration.

## INTRODUCTION

The Khapra beetle, *Trogoderma granarium* Everts (Coleoptera: Dermestidae), is an important and destructive insect pest of stored products (Naseri and Borzoui, 2016). This pest causes economic losses, particularly in tropical and subtropical regions of Asia and Africa (Burges, 2008). Infestation of seeds and food commodities by *T. granarium* larvae and their cast skins and hairs cause loss of biomass and food quality of stored products (Jood et al., 1996). In response to food shortage and unfavorable conditions (i.e. temperatures below 30 °C), the larvae enter diapause, remain relatively inactive, and rarely feed. Diapausing larvae tend to leave the food and aggregate in crevices of the buildings (Burge, 1962).

The best known and extensively researched mechanisms of insect cold hardiness are carbohydrate cryoprotectants, AFPs, and INAs or INPs. All contribute protective mechanisms that deal with problems of water and ice at subzero temperatures and all have been extensively reviewed in the past including two books (Lee and Denlinger, 1991; Denlinger and Lee, 2010) and many review articles (some examples: Storey and Storey, 1988; 1991; Zachariassen and Kristiansen, 2000; Duman, 2001; Davies et al., 2002; Block, 2003; Graham et al., 2007; Clark and Worland, 2008; Doucet et al., 2009). Briefly, both freeze-avoiding and freeze-tolerant species accumulate polyol cryoprotectants; in freeze-avoiding species, polyols permit colligative suppression of supercooling point to prevent body freezing, though in freeze-tolerant species polyols offer protection against intracellular freezing when ice accumulates in extracellular compartments. In cryoprotective dehydration, the disaccharide trehalose plays the major role in macromolecular protection. AFPs are critical to supercooling point suppression in freeze-avoiding species but can also be found in freeze-tolerant species where they appear to function in inhibition of ice recrystallization. Interestingly, new research has made a novel discovery of a new class of biological antifreeze—xylomannan glycolipids (Walters et al., 2009). Walters et al. (2011) demonstrated their presence and antifreeze action in six insect species, as well as in one plant and one frog species; much characterization of their action remains to be done. Ice nucleators act in freeze-tolerant species to trigger ice formation at high subzero temperatures and manage ice growth in extracellular compartments; insect INPs are often lipoproteins (Trautsch et al., 2011).

The AMP-activated protein kinase (AMPK) is proving to be a major regulator of catabolic vs. anabolic poise in cells, its actions favoring the former and inhibiting the latter. AMPK was first discovered as a protein kinase that was allosterically activated by AMP accumulation under low energy conditions (e.g., hypoxia) and it is often called the energy sensor or the fuel gauge of the cell (Hardie, 2007; Hue and Rider, 2007). Also, AMPK is now known to respond to various upstream kinases that allow other signals to have input on AMPK targets. The best-known action of AMPK is phosphorylation and inactivation of acetyl-CoA carboxylase (ACC), which inhibits lipogenesis and promotes fatty acid oxidation under energy-limiting conditions. AMPK activation also exerts inhibitory control over carbohydrate storage (by inhibiting glycogen synthase) and protein synthesis (by activating the protein kinase that inactivates the ribosomal eukaryotic elongation factor-2 (eEF2). A series of recent studies have consistently shown AMPK activation in animals transitioning into hypometabolic states (e.g., frog freeze tolerance, turtle and fish anaerobiosis, nematode dauer) (summarized in Rider et al., 2011).

In our study, we hypothesize that variation in cold tolerance in *T. granarium* larvae acclimated with low temperature arises from variation in the ability to change in AMP-activated protein kinase and protein phosphatases activity, maintain ion balance and increases the concentration of cryoprotectant contents in the cold. We thus predict that if acclimated with the low-temperature conditions, cold tolerant Drosophila species would: 1) the enzyme activities varies according to the thermal regimes, 2) better maintain [Na+] and [K+] balance in their hemolymph fluid, and 3) changes in enzyme activity and ion balance increases the concentration of cryoprotectant contents, and improves cold tolerance.

## MATERIALS AND METHODS

### Chemicals

All chemicals used for analysis were purchased from Sigma-Aldrich (St. Louis, MO, USA).

### Insect rearing

The *Trogoderma granarium* population used for the experiments was obtained from cultures that had been originated from stored rice seeds from Karaj (Iran) and maintained for 2 years in the Laboratory of Entomology, Vali-e-Asr University of Rafsanjan, Rafsanjan, Iran. The insects were fed on broken wheat seeds (*Triticum aestivum* L.) under a controlled environmental chamber at 33 ± 1 ºC with 65 ± 5% RH (by using saturated salt solution) and a photoperiod of 14:10 (L:D) h, as described by Nouri-Ganbalani and Borzoui (2017).

### Acclimation treatments

Beetles were raised from egg to the fourth instar in translucent plastic containers (diameter 15 cm, depth 6 cm) with a hole covered by a 50 mesh net for ventilation, containing broken wheat seeds. *T. granarium* 4^th^ instar larvae were divided into four groups: warm-acclimated as control (WA), cold-acclimated (CA), fluctuating-acclimated (FA) and rapid cold-hardened (RCH). For WA treatment, one hundred individuals were put in a translucent plastic cups containing food and kept in standard rearing conditions. For CA treatment, one hundred individuals were put in a translucent plastic cups containing food, cooled in a programmable refrigerator from rearing conditions to 15 ºC at a rate of 0.5 °C min^−1^ and kept at this temperature at 65 ± 5% RH with a 14:10 h (L:D) light cycle for the 10 days. Thereafter, the temperature was lowered to 5 ºC at the same rate and the larvae were kept at this temperature at 65 ± 5% RH with a 14:10 h (L:D) light cycle for the 10 days. For FA treatment, one hundred individuals were put in a translucent plastic cups containing food, cooled in a programmable refrigerator from rearing conditions as explained in the cycle: 240 min at 5 °C followed by 20 min at -10 °C followed by 240 min at 5 °C followed by 940 min at 33 °C, at 65 ± 5% RH with a 14:10 h (L:D) light cycle. This cycle repeated 10 consecutive days. For RCH treatment, the larvae were transferred from their rearing conditions to a programmable refrigerator at 0 ºC for four hours. After the treatment period, survived larvae were used for subsequent experiments. The larvae able to walk were counted as alive, and larvae that were either not showing any movement in their appendages or moving but unable to walk were counted as dead.

### Enzymes preparation and assay

The whole body of acclimated larvae of *T. granarium* was used; it was not feasible to separate out individual tissues. For all enzymes, activities were expressed as Unit per gram wet mass. Assays were repeated five times.

#### APK

Individuals were rapidly weighed and homogenized 1:10 (w/v), with a few crystals of phenylmethylsulfonyl fluoride (PMSF) added, using a pre-cooled homogenizer (Teflon pestle) in ice-cold potassium phosphate buffer (20mM; pH 6.8), X-mercaptoethanol (15mM) and EDTA (2mM). The homogenates were centrifuged at 13000 *g* for 3 min (5 °C). Following centrifugation, the supernatant was pooled and stored on ice for subsequent use. The activity of APK was assayed by the procedure of Pfister and Storey (2002a). In brief, ^32^P from ^32^P-ATP was incorporated onto Kemptide (LRRASLG), a synthetic phosphate-accepting peptide, in the presence of 0.1mM adenosine 30,50-cyclic monophosphate. One unit of APK activity is defined as the amount of enzyme required to catalyze the incorporation of 1 nmol ^32^P onto the substrate per minute at 23 °C.

#### PP1

Individuals were rapidly weighed and homogenized 1:3 (w/v) using a pre-cooled homogenizer (Teflon pestle) in ice-cold Buffer A [Tris–HCl (20 mM; pH 7.4), EDTA (2 mM), EGTA (2 mM), X-mercaptoethanol (15 mM)] containing the protease inhibitors: PMSF (1 mM), TPCK (0.1 mM), aprotinin (1 mg/ml) and benzamidine (5 mM). The homogenates were centrifuged at 1000 *g* for 3 min (5 °C). Following centrifugation, the supernatant was carefully collected and assayed immediately for active PP1. Estimates of PP1 activities at physiological levels of modulating proteins and other factors done based on assays of concentrated extracts (Toth et al., 1988). PP1 activity was estimated at 23 °C by monitoring ^32^P cleavage from ^32^P-labeled phosphorylase (Pfister and Storey, 2002b). One unit of PP1 activity is defined as the amount of enzyme required to releases 1 nmol of phosphate per minute at 23 °C.

#### PP2

Individuals were extracted as for PP1 except for a 1:10 (w/v) dilution. The homogenates were centrifuged at 13000 *g* for 20min (5 °C). Following centrifugation, the supernatant was carefully collected and desalted by centrifugation at low-speed for 1min (at room temperature) through 5 ml Sephadex G-25 columns equilibrated in ice-cold Buffer A. The eluant was collected, passed through a second, fresh column and stored on ice for subsequent use. The activity of PP2A and PP2C were assayed by the procedure of Cowan et al. (2000). PP2A activity was measured as the difference in activity in the presence (blank) versus absence of okadaic acid (2.5 nM). To assess the PP2A activity, the reaction mixture containing peptide RRA(pT)VA (150 mM), EGTA (0.2 mM), X-mercaptoethanol (0.02%), and imidazole (50 mM), pH 7.2, and 10 µl of enzyme extract was incubated for 40 min. The reaction was terminated by adding 50 ml of malachite green dye solution [ammonium molybdate (10%) and malachite green dye (2%), both in HCl (4 N) mixed 1:3 v/v and diluted 2:3 v/v with distilled, deionized water, Tween 20 (0.05%) and Triton-X-100(0.05%)] (Ekman and Jaeger, 1993). Reactions were run in 96-well microplates and the absorbance was read at 595 nm. Appropriate blanks, which TCA had been added prior to the substrate, were prepared for each treatment. The activity of PP2C was assayed at the same except for the presence of okadaic acid (2.5 nM) and incubation of the reaction mixture for 90 min; PP2C was detected as the difference in activity in the absence versus presence of MgCl_2_ (10 mM) (Cowan et al., 2000).

### Ion concentration

Ion concentrations were measured in the hemolymph (*n* = 5) as previously described by MacMillan et al. (2015) with some modification. [K+] and [Na+] were measured in the hemolymph at 0, 1, 2, 4, 6 and 12 h after exposure to -10 °C in individuals treated with different thermal regimes (100 individuals from each treatment and time point). Hemolymph was sampled (and weighed, to estimate volume) using a micropipette from an incision made at the coxal joint of a hind leg while applying gentle pressure to the abdomen to allow the hemolymph to flow into the tube. Then, the hemolymph was transferred to a 0.5 ml Eppendorf tube, which was placed in a microcentrifuge (DW-41-230, Radiometer A/S, Brønshøj, Denmark) and spun for 15 s to separate hemolymph from debris. Afterward, 1–5 μl sample of hemolymph was transferred by pipette to a 2 ml buffer solution containing 100 ppm lithium salt. After the preparation described above, the [Na+] and [K+] was measured from the hemolymph using an atomic absorption spectrometry (AAS; Model iCE 3300, Thermo Scientific, Waltham MA, USA) and comparisons to standard curves.

### Whole-body glycogen and sugar alcohols quantification

The whole-body glycogen and polyol profiles of acclimated larvae of *T. granarium* were repeated with five replicates (one individual from each treatment and time point) for each experiment at the end of the thermal regimes. All concentrations are expressed as microgram per gram of dry weight.

#### Glycogen

The glycogen content of the larvae was estimated using modified anthrone method as described by Heydari and Izadi (2014). The larvae were weighed and homogenized in 200 μl of 2% Na_2_SO_4_. Thereafter, 1300 μl chloroform-methanol (1:2) was added to the homogenate. The homogenates were centrifuged for 10 min at 7150 *g* and the supernatant was removed. The pellet was washed in 400 μl of 80% methanol and 250 μl distilled water was added before the heating for 5 min at 70 °C. Subsequently, 200 μl of the solution was incubated with 1 ml of anthrone for 10 min at 90 °C. After cooling at room temperature, the absorbance of the solution was measured at 630 nm. The glycogen content was determined by comparison to a standard curve that was prepared using glycogen.

#### Sugar alcohols

The extraction, derivatization, and analytical procedures (gas chromatography coupled to mass spectrometry) were similar to those described by Heydari and Izadi (2014). After the homogenization of individual larvae in 1.5 ml of 80% ethanol and centrifugation (twice repeated), the supernatant (20 ml) was run along with standards of polyols from 1500 to 5500 ppm. Trehalose, sorbitol, myo-inositol and glucose were analyzed by HPLC (Knauer, Berlin, Germany) using a carbohydrate column with 4 μm particle size (250 mm × 4.6 mm, I.D., Waters, Ireland).

### Determination of supercooling point

The supercooling point (SCP) was determined for the acclimated larvae of *T. granarium* (*n* = 15). To determine SCP, individual larvae were placed on a thermocouple (NiCr–Ni probe) connected to an automatic temperature recorder (Testo 177-T4, Testo, Germany) within a programmable refrigerated test chamber. The temperature of the refrigerated test chamber was reduced from experimental conditions to -30 °C, at a rate of 0.5 °C min^−1^. The lowest temperature reached before an exothermic event that occurred caused by the release of latent heat was taken as the SCP of the individual (Mohammadzadeh and Izadi, 2016).

### Cold-tolerance assays

In total two separate experiments were done to study cold tolerance: (1) an acute cold-tolerance assay at subzero temperature for 1 h, and (2) an experiment measuring cold-tolerance at -5, -10, - 15 and -20 for 24 h. Five replicates and 15 larvae for each replicate were used at each treatment and temperature point. To estimate the acute cold exposure, the larvae treated with different thermal regimes were exposed to acute low temperature (-5 °C to -20 °C). Survival rate was assessed after 1 h. Finally, LT_80-1 h_ was calculated as the lowest temperature at which 80% of the larvae died after 1 h exposure (Sinclair and Rajamohan, 2008). To estimate the cold tolerance, the larvae treated with different thermal regimes were kept in a programmable refrigerated test chamber, whose temperature was lowered slowly (0.5°C min^−1^) from experimental conditions to the desired treatment temperature (-5, -10, -15 and -20 ± 0.5°C) and held at each temperature for 24 h. The mortality of larvae was recorded via direct observation. The larvae showing no movement in their appendages were judged to be dead (Mohammadzadeh and Izadi, 2016).

### Statistical analysis

Data were initially tested for normality (Kolmogorov−Smirnov test) and homoscedasticity (Levene’s test) before subjecting them to ANOVA. All the data were analyzed using SAS ver.9.2 program (PROC GLM; SAS Institute 2009). Statistical analyses were performed, based on a completely randomized design, using one-way analysis of variance (ANOVA) followed by a post hoc Tukey’s test at α = 0.05.

## RESULTS

### Effect of thermal regimes on enzymes activity

Profiles of enzymes activities in *T. granarium* under different thermal regimes are shown in Figure 1. Based on the results of this study, the highest and lowest activities of APK (55 and 25 units/gwm, respectively) and PP2C (27 and 14 units/gwm, respectively) were observed at CA and FTT treatments, respectively. No significant differences were observed in the activities of these two enzymes between control and RCH. If the activities of these two enzymes changed in the same norm in different regimes, but APK was found to be much more active than PP2C. In control, the activity of APK was about 33 units/gwm, whereas the activity of PP2C was about 19 units/gwm. The activity of both enzymes increased by CA and reached to the highest levels of 55 and 27 units/gwm, respectively. The activity of PP1 and PP2A also showed more or less of the same norm under different regimes. These norms were completely different from those of the two other enzymes. The highest (28 units/gwm) and lowest (12 units/gwm) activities of PP1 were observed at control and CA treatments, respectively. In case of PP2A, the highest level of activity was recorded for control and FTT treatments (1.1 and 1.2 units/gwm, respectively), whereas the lowest level of activity was shown in CA and RCH (0.5 and 0.6 units/gwm, respectively). In PP1, the highest (28 units/gwm) and lowest (13 units/gwm) levels of activity were observed in control and CA regimes, respectively.

**Figure1.**
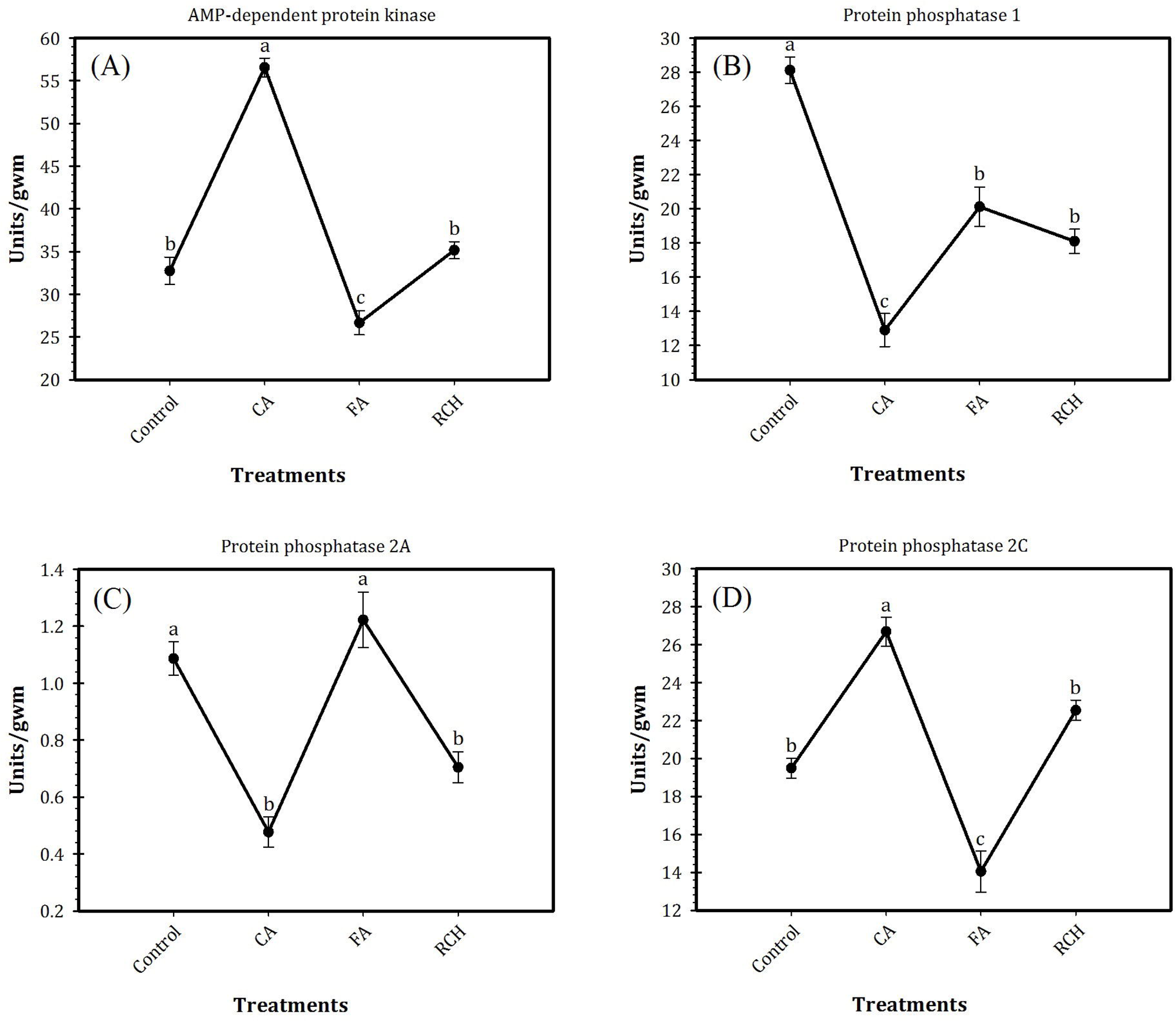
Profiles of AMP-dependent protein kinase and protein phosphatase activities in *Trogoderma granarium* 4^th^ instar larvae following different thermal regimes. (A) total PKA, (B) total PP1, (C) PP2A, (D) PP2C. Each point is average of five replications. The means followed by different letters are significantly different (Turkey’s test, *P* < 0.05).

### Effects of thermal regimes on hemolymph Na^+^ and K^+^ concentrations

Our results showed that in all the regimes as well as in the control, the concentration of Na^+^ decreased, whereas the concentration of K^+^ increased with an increase in exposure time of the larvae at -10 ºC (Fig 2). In all the regimes, at the beginning time of exposure (0 h at -10 ºC) concentration of Na^+^ was about 70 mM. The concentration of Na^+^ increased and reached to the highest level after 1 h exposure at -10 ºC. Then, the concentration of this ion in control and different thermal regimes steady state decreased with increasing exposure time. The sodium/potassium ratio decreased with an increase in exposure time. During different times of exposure at -10 ºC, the highest and lowest levels of Na^+^ and K^+^ were recorded for control and CA treatment, respectively. In addition, at the highest levels, the concentration of Na^+^ was about four times more that of K^+^.

**Figure 2.**
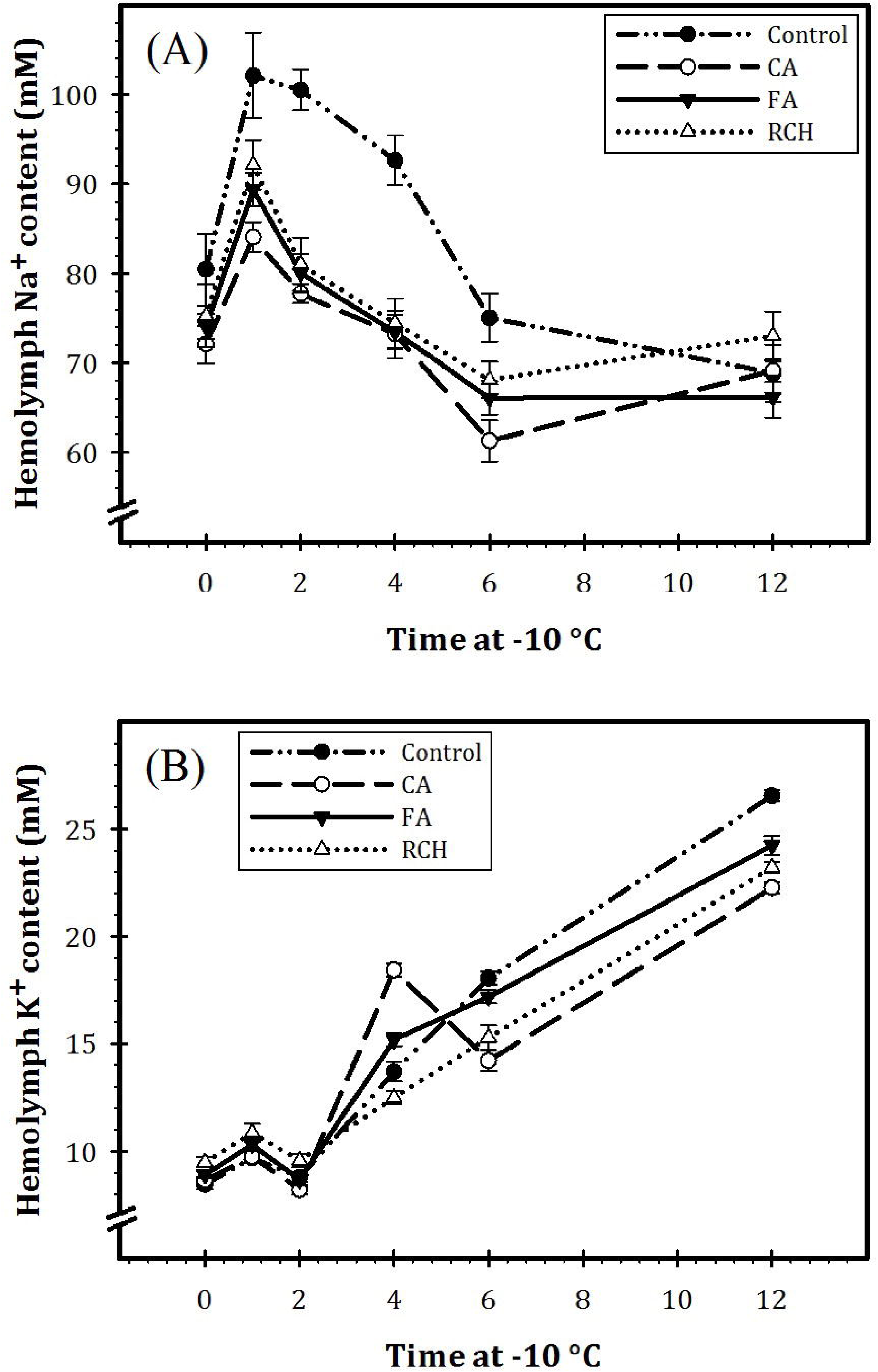
Hemolymph Na^+^ (A) and K^+^ (B) concentrations before and immediately following 1, 2, 4, 6 and 12 h at 10 °C in *Trogoderma granarium* 4^th^ instar larvae following different thermal regimes. Each point is average of five replications. The means followed by different letters are significantly different (Turkey’s test, *P* < 0.05).

### Effects of thermal regimes on carbohydrate contents

Glycogen content in control larvae with 34.4 µg/gdw was at the highest level and reached the lowest level of 23.0 in CA larvae. No significant differences were observed between glycogen contents in FTT, RCH, and control (Table 1). Trehalose and myo-inositol were found to be the predominant low molecular weight of carbohydrates in all the regimes. Changes in low molecular weight carbohydrate contents were reversely proportional to change in glycogen content. The highest and lowest contents of trehalose and myo-inositol were observed in CA (16.5 µg/gdw) and control (9.9 µg/gdw), respectively. No significant differences were observed in sorbitol contents between control and different thermal regimes. The highest and lowest glucose contents were observed in control (0.49 µg/gdw) and CA (0.14 µg/gdw), respectively.

**Table 1.**
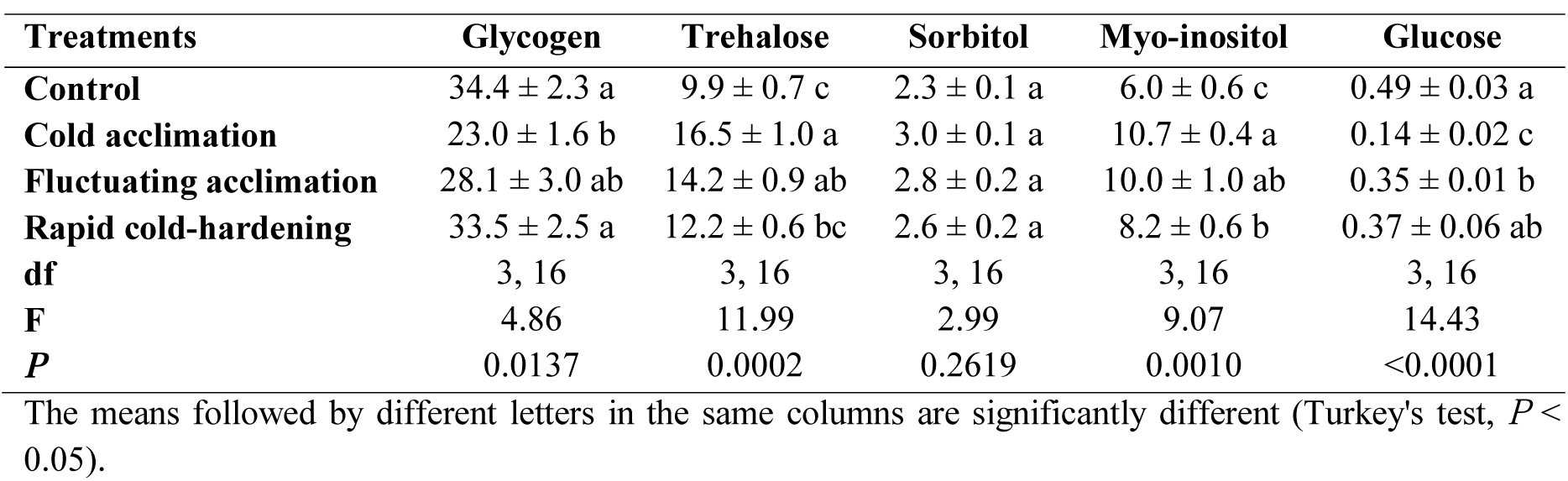
Glycogen and carbohydrate contents (*n* = 5) of *Trogoderma granarium* 4^th^ instar larvae following different thermal regimes.

| Treatments | Glycogen | Trehalose | Sorbitol | Myo-inositol | Glucose |
| --- | --- | --- | --- | --- | --- |
| **Control** | 34.4 ± 2.3 a | 9.9 ± 0.7 c | 2.3 ± 0.1 a | 6.0 ± 0.6 c | 0.49 ± 0.03 a |
| **Cold acclimation** | 23.0 ± 1.6 b | 16.5 ± 1.0 a | 3.0 ± 0.1 a | 10.7 ± 0.4 a | 0.14 ± 0.02 c |
| **Fluctuating acclimation** | 28.1 ± 3.0 ab | 14.2 ± 0.9 ab | 2.8 ± 0.2 a | 10.0 ± 1.0 ab | 0.35 ± 0.01 b |
| **Rapid cold-hardening** | 33.5 ± 2.5 a | 12.2 ± 0.6 bc | 2.6 ± 0.2 a | 8.2 ± 0.6 b | 0.37 ± 0.06 ab |
| **df** | 3, 16 | 3, 16 | 3, 16 | 3, 16 | 3, 16 |
| **F** | 4.86 | 11.99 | 2.99 | 9.07 | 14.43 |
| **P** | 0.0137 | 0.0002 | 0.2619 | 0.0010 | <0.0001 |

The means followed by different letters in the same columns are significantly different (Turkey's test, $P < 0.05$ ).

### Effect of thermal regimes on SCP and survival of the larvae

Data in table 2 showed that under CA and FTT thermal regimes supercooling points of the larvae decreased to the lowest level (about -22 ºC), which significantly lower than SCPs of control and RCH regimes. No significant difference was observed between SCPs of control and RCH.

**Table 2.**
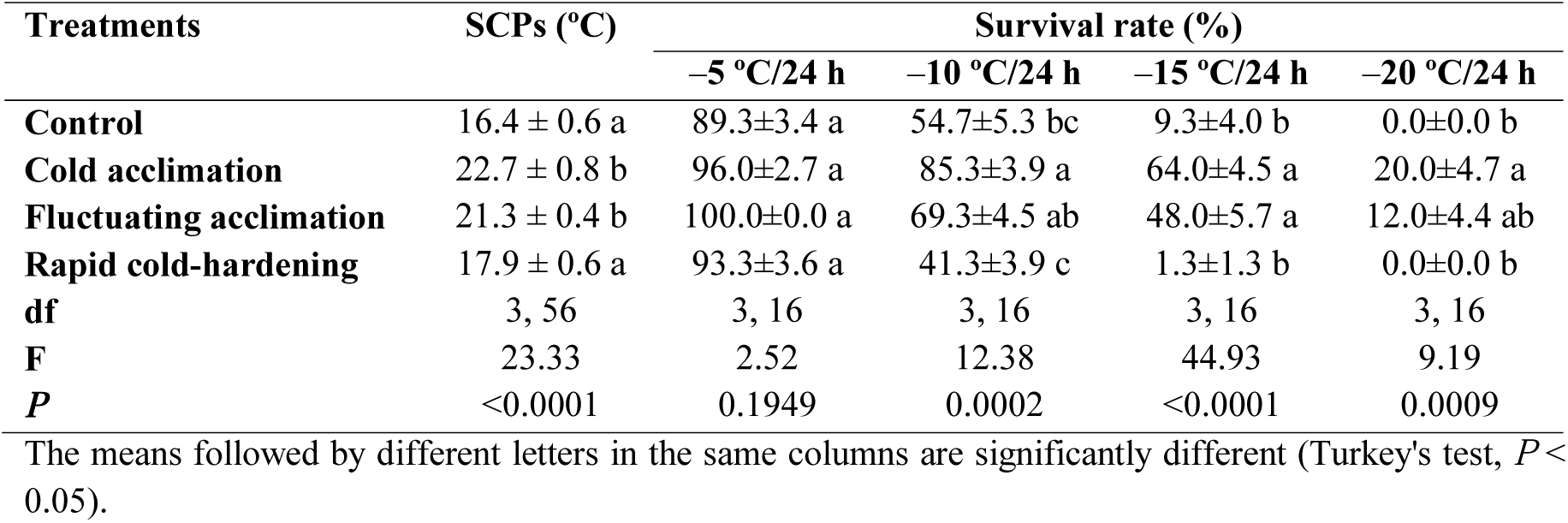
Relationship between low temperature survival rate (*n* = 5) and supercooling points (*n* = 15) of *Trogoderma granarium* 4^th^ instar larvae following different thermal regimes.

Our results also showed a profound effect of CA on the LT_80_ value of the larvae (Fig 3). LT_80_s of the larvae at control, RCH, FTT and CA regimes were calculated as 11, -14, -19 and -21 ºC, respectively. In CA larvae, the temperature required for 80% mortality decreased about 10 ºC compared with control larvae. In addition, the temperature necessary for the beginning of mortality in CA regime was 10 ºC lower than that of control. In all the regimes, the survival rate of the larvae decreased with a decrease in the temperature and increase in the exposure time. Cold acclimated larvae showed the highest cold hardiness in the -15 and -20 ºC. In control, RCH, FTT, and CA larval mortality was begun at -20, -25, -27 and -30 ºC, respectively.

**Figure 3.**
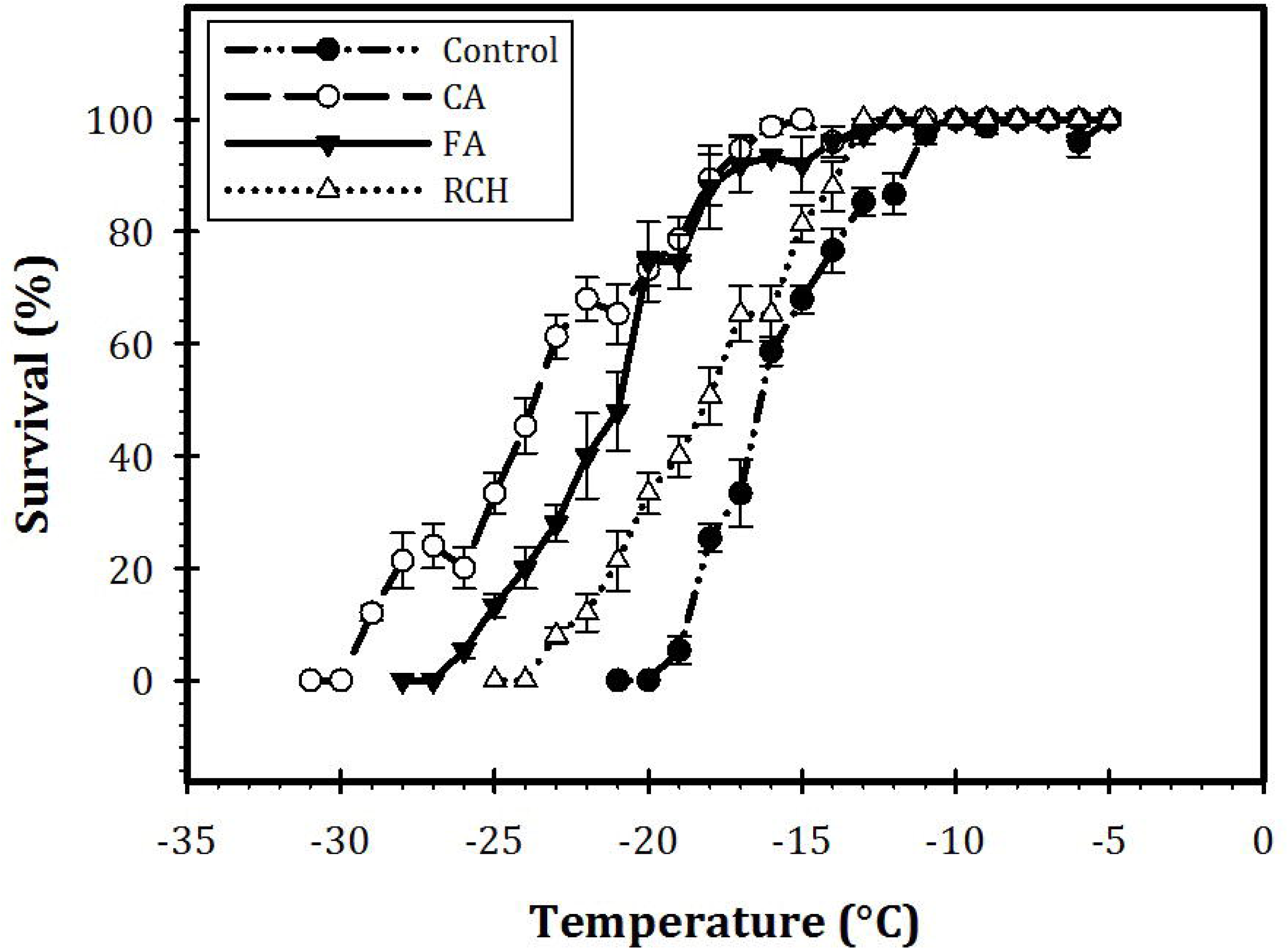
Survival of *Trogoderma granarium* 4^th^ instar larvae following different thermal regimes after acute low-temperature exposure. The dotted line indicates 80% mortality (Lt_80–1 h_).

## DISCUSSION

The APK as a downstream component of a kinase cascade has several functions (regulation of glycogen, sugar, and lipid metabolism) and cellular targets. This enzyme can be regulated by hormones and cytokines (Hardie, 2010). Our findings showed that APK is a predominant signal transduction enzyme of *T. granarium* last instar larvae. Activities of the tested enzymes could be rated as APK>PP1>PP2C>PP2A. The results of previous studies strongly support the findings of the current study. Pfister and Storey (2006a) demonstrated changes in the activities of APK, PP1, PP2A and PP2C in a freeze-avoiding insect, *Epiblema scudderiana* (Clemens) (Lep.: Olethreutidae) in winter and during exposures of the larvae to subzero temperatures. They demonstrated a limited change and the role for APK in overwintering larvae, but the activities of PP2A and PP2C increased when larvae were exposed to -20 degrees ºC. In another study, Pfister and Storey (2006b) studied changes in the activities of the same enzymes in the goldenrod gall fly, *Eurosta solidaginis* Fitch (Dip.: Tephritidae), and resulted in an increase in APK and a decrease in PP1 activity over the winter season and/or at subzero temperature.

The findings of the present research revealed that the shape of the thermal reaction curve in APK and PP2C follows the same norm that is different from those of PP1 and PP2A. PP2A and 2C showed opposite trends in activity and different thermal reaction norms. Indeed, thermal fluctuation caused the highest changes in the enzyme activities, whereas larvae of RCH treatment showed the lowest level of the enzyme activities.

Our previous study has shown that larvae of *T. granarium* are freeze-avoidant or freeze-intolerant (Mohammadzadeh and Izadi, 2017). In these freeze-intolerant larvae, several metabolic adaptations, including synthesis of polyols and low molecular weight carbohydrates (as cryoprotectant) have been developed for survival at subzero temperatures and harsh environmental conditions. The results of the current study indicated a significant enhancement in larval survival and cold tolerance under CA regime. The results of our study also revealed the profound impact of CA on carbohydrate contents of the larvae. In the CA larvae, cold hardiness of the larvae was at the highest value and major cryoprotectants such as trehalose and myo-inositol were at the highest levels, but glycogen reached the lowest concentration. So, it could be concluded from these results that cold acclimation is important in the conversion of glycogen to low molecular weight carbohydrates, which act as a cryoprotectant to enhance cold hardiness of the larvae. High level of cryoprotectants (e.g. trehalose) is essential for reduction of supercooling in freeze avoidant species or to prevent intracellular ice formation in freeze tolerant insects.

The induction of insect cold hardiness and related adaptations require the intermediary action of signal transduction enzymes (Pfister and Storey, 2006b). In agreement with the above, in the current study, the activity of APK, as a signal transduction enzyme, in the cold-acclimated larvae increased and reached to the highest level. An increase in the activity of this enzyme was coincident with the increase of cold hardiness, cryoprotectants concentration, and survival rates. These findings strongly support a regulatory role for APK and PP1 in the synthesis of cryoprotectants from glycogen. In the cold-acclimated larvae, glycogen content decreased with the increase in trehalose content and APK activity. So, it is reasonable to conclude that APK may be responsible for shutting down glycogen synthesis and activates conversion of glycogen to trehalose. For cryoprotectants synthesis, conversion of glycogen to polyols or sugar alcohols is necessary (Storey and Storey, 2012). In this process, the role of PP2C is more limited than APK and there is no role for PP1 and PP2A. In overwintering larvae of *E. scudderiana*, PP1 was found to be responsible for shutting off glycogenolysis, whereas a limited role was attributed to PKA. In this moth, the activity of PP2A and PP2C increased by exposing the larvae to -20 ºC (Pfister and Storey, 2006a). A recent years study suggests a big role for APK in insect cold hardiness and diapause. Rider et al. (2011) resulted in 2-fold higher activity of APK in winter larvae of *E. solidaginis* and *E. scudderiana* in compared to summer ones. Joanisse and Storey (1995) found an increase in activities of glycogenolytic and hexose monophosphate shunt enzymes in cold-acclimated *E. scudderiana* larvae, which resulted in the conversion of glycogen into glycerol as a cryoprotectant. In *E. solidaginis* cold acclimated larvae, an increase in activity of glycogen phosphorylase with a decrease in activity of glycolytic enzymes may be responsible for the temperature-dependent switch from glycerol to sorbitol synthesis.

To overcome adverse effects of low temperature, several physiological mechanisms have been developed in insects of cold and temperate zones. Three main groups of these mechanisms are: 1-physiological, biochemical and metabolic alterations (cryoprotectant synthesis and synthesis of antifreeze proteins and/or ice nucleating agents), 2-change in cell function (modification of membranes, regulation of ion-homeostasis and mobilization of cryoprotectants) and 3-alternation in gene expression (up-regulation of stress-related genes) (Overgaard et al., 2005; Overgaard et al., 2007; Lee and Denlinger, 2010; Teets and Denlinger, 2013; Overgaard et al., 2014).

In our study, trehalose content was at the highest level in cold-acclimated larvae. Trehalose, as a cryoprotectant, contributes to stabilize the lipid bilayer of the cell membrane (Crowe et al., 1992; Storey and Storey, 2012). The results of this study are in agreement with the results of Pfister and Storey (2006b). They showed several metabolic adaptations in freeze-tolerant larvae of the goldenrod gall fly for subzero survival. Several other studies reported this sugar as a cryoprotectant in cold hardy insect species (Behroozi et al., 2012; Sadeghi et al., 2012; Bemani et al., 2012; Mohammadzadeh and Izadi, 2017b; Heydari and Izadi, 2014). In disagreement with our results, Mollaei et al. (2016) resulted that high cold tolerance of larvae of *Kermania pistaciella* Amsel (Lep.: Oinophylidae) was not associated with accumulation of cryoprotectants during overwintering.

Based on the results of the current study, the SCP reduced to the lowest level in cold-acclimated larvae. The decrease in SCP was proportional to an increase in cold hardiness of the larvae which in turn was associated with an increase in the enzyme activities and cryoprotectant accumulations. Thus, a strong relation between enhanced cold hardiness, elevated enzyme activities, accumulated cryoprotectants and decreased SCP of the Khapra beetle cold acclimated larvae can be concluded from our results. On the other hand, in the cold-acclimated larvae, enhancement of cold hardiness is a function of both a decrease in SCP and an increase in trehalose synthesis and accumulation. Hiiesaar et al. (2001) resulted that the mean supercooling point of *Leptinotarsadecemlineata* (Say) (Col.: Chrysomelidae) decreased from -10.5 in non-acclimated to -17.5 ºC in cold-acclimated adult beetles. In prepupae of *Hermetia illucens* (L.) (Dip.: Stratiomyidae), the SCP was unaffected by cold acclimation, but cold hardiness increased in compared to control (Spranghers et al., 2017).

Insects have also the capability to improve cold tolerance and survival rate in a short period, called rapid cold-hardening (RCH) (Teets and Denlinger, 2013). In the current study, RCH had no significant effects on enzyme activities, survival rates, and cryoprotectant accumulation. This correlates well with the previous studies. Overgaard et al. (2014) in adults of *Drosophila melanogaster* Meigen (Dip.: Drosophilidae) reported no effect of rapid cold hardening (RCH) on the activity of glycogen phosphorylase. They found a small increase in glucose content, whereas, trehalose content remained unchanged following RCH. Kelty and Lee (2001) in rapidly cold-hardened adults of *D. melanogaster* found no change in the levels of Hsp70 and carbohydrate cryoprotectants. In disagreement with our results, Lee et al. (2006a) reported that RCH significantly increased survival of *Belgica antarctica* Jacobs (Dip.: Chironomidae) larvae. Lee et al. (2006b) determined that RCH increased membrane fluidity of fat body cells of *Sarcophaga bullata* (Parker) (Dip.: Cyclorrhapha) adult flies. They suggested that “membrane characteristics may be modified very rapidly to protect cells against cold-shock injury”. In adults of *Thrips palmi* Karny (Thysan.: Thripidae) RCH caused accumulation of cryoprotectants mainly trehalose (Park et al., 2014). So, based on our results and results of other researchers, it is reasonable to conclude that insects may become cold hardy by RCH if RCH participates in the accumulation of cryoprotectants (polyols and sugar alcohols). In our study, no accumulation of cryoprotectants and consequently no cold hardiness were determined in RCH regime. Jakobs et al. (2017) found no evidence that acute cold tolerance of *D. suzukii* larvae could be improved by RCH.

Chilling insect at low temperature causes a loss of extracellular ions and water homeostasis (MacMillan et al., 2015). In a recent year experiment, MacMillan et al. (2015) examined the capacity of chill susceptible *Drosophila* species malpighian tubules (MT) and demonstrated that MT lost [Na^+^] and [K^+^] selectivity at low temperatures, which participate in a loss of Na^+^ and water balance and an increase in extracellular [K^+^]. These findings strongly support the results of the current study. Based on our results, exposure of the larvae at low temperature caused a substantial decrease in [Na^+^] and an increase in [K^+^]. This finding strongly supports the concept that low temperature reduced [Na^+^] and [K^+^] selectivity of MT, which contributed to a decrease in [Na^+^], a harmful increase in [K^+^] and consequently, an accumulation in chill injuries. Insect cold hardiness is strongly associated with the ability of MT to retain ions (Particularly K^+^ and Na^+^) and water balance during cold exposure (MacMillan et al., 2015; Andersen et al., 2017). MacMillan et al. (2015) concluded that chill tolerant *Drosophila* species maintained K^+^ secretion better than chill susceptible species and suppressed K^+^ reabsorption during cold exposure. These data, therefore, are strongly in agreement with our findings. As our results show, at the beginning of the cold exposure, hemolymph [Na^+^] is at the highest level, whereas; [K^+^] is at the lowest level. By increasing exposure time, [Na^+^] decreased and [K^+^] increased. In most insects, hemolymph [Na^+^] is significantly higher than [K^+^]. So, Na^+^ ions tend to leak into the gut lumen while K^+^ ions tend to leak into the hemolymph. At normal temperature, passive ion movements from the MT lumen to the hemolymph and from the hemolymph to the MT lumen are regulated by the energy-demanding proton pump located at the apical membrane MT epithelial cells. At low temperature, [Na^+^] leak away from the hemolymph to the MT lumen and so, its concentration reduces in hemolymph leading to increasing in [K^+^] in the hemolymph. Increase in hemolymph [K^+^] causes depolarization of cell resting potential and this depolarization may be a primary reason of cold-induced injury (Kostal et al., 2004; MacMillan et al., 2014; MacMillan et al., 2015; Andersen et al., 2017). We can conclude from our results that the gene expression of ion homeostasis and consequently water balance has been altered cold acclimated larvae of the Khapra beetle. The same results have been reported by Gerken et al. (2015); MacMillan et al. (2015a) and Des Marteaux, L. E. (2017).

## ACKNOWLEDGEMENTS

The authors thank the Vali-e-Asr University of Rafsanjan (Rafsanjan, Iran), for cooperation by support for the experiment.

## AUTHOR CONTRIBUTIONS

MM and HI conceived and designed research, conducted experiments, contributed analytical tools and analyzed data, and wrote the manuscript.

